# Distribution, prevalence, and causative agents of fungal keratitis: A systematic review and meta-analysis (1990 to 2020)

**DOI:** 10.1101/2021.04.19.440430

**Authors:** Kazem Ahmadikia, Sanaz Aghaei Gharehbolagh, Bahareh Fallah, Mahsa Naeimi, Pooneh Malekifar, Saeedeh Rahsepar, Muhammad Ibrahim Getso, Savitri Sharma, Shahram Mahmoudi

## Abstract

Fungal keratitis is a sight-threatening infection with global distribution. In this systematic review and meta-analysis we collected all the articles with data on the prevalence of fungal keratitis among various patient groups from January 1, 1990 to May 27, 2020. The 169 eligible articles were divided into 6 groups. The pooled prevalence was variable with values ranging from 0.05% among post-keratoplasty patients to 43.01% among patients with a clinical suspicion of fungal keratitis. Except for post-keratoplasty cases (yeast: 51.80%), in all patient groups moulds were more common than yeasts. Although more than 50 distinct species of fungi have been found to cause fungal keratitis, *Fusarium* species followed by *Aspergillus* species were the most common causes of the disease. In general, 9.29% (95% CI 6.52, 12.38) of fungal keratitis cases were mixed with bacterial agents.

## 1. Introduction

Keratitis describes a group of acute or chronic inflammatory disorders occurring in the cornea following any factors disrupting the protective mechanism of the outer layer of the eye [1]. The inflammation may be of allergic (reactive), physical, chemical or infective (bacteria, fungi, parasites and viruses) origin [1]. Infectious or microbial keratitis, one of the most serious eye infections, has long been acknowledged as the leading cause of visual impairment and blindness worldwide [2]. Very appropriately, infectious keratitis is now included among neglected tropical diseases by world health organization [3]. Due to the non-specific signs and symptoms, and rapid progression, microbial keratitis is diagnostically challenging for physicians [4].

The prevalence rate and epidemiological distribution of fungal keratitis (FK) are strongly associated with geographical locations and widely vary throughout the world, even between different regions of the same country and in different groups of individuals [5]. FK is globally getting increasing attention particularly in developing countries, tropical and subtropical regions [6, 7] where approximately half of the world’s fungal keratitis cases belong [8], therefore, the contribution of FK, as one of the major causes of visual loss cannot be neglected [9].

Characteristic clinical features of FK have been described, however, they are not pathognomonic enough and often mimic (masquerades) bacterial or parasitic keratitis [2]. Therefore, in absence of laboratory diagnosis majority of the cases may be treated empirically [2, 10], resulting in poor outcome that may progress to endophthalmitis, particularly if left untreated [5, 11, 12]. Emerging fungal pathogens and resistance to existing antifungal agents have further contributed to poor prognosis in FK [5, 6].

A healthy cornea is rarely infected with fungal agents [13]. Traditionally, FK is considered to be a disease prevalent in rural settings occurring in traumatized eyes by vegetative materials or soil-contaminated objects in middle-aged agriculturists and labourers. These are evidenced to be the most susceptible eyes for fungal infection of the cornea in low-income countries [1, 14]. Conversely, contact lens usage (CLU) is the primary culprit, predisposing hosts to FK in developed countries [1, 7, 14].

The most prevailing species implicated in FK, include those belonging to *Aspergillus, Fusarium, Candida, Curvularia*, and *Penicillium* genera in descending frequency [15–19]. Most of these species are environmental residents which invade traumatized or immunologically weakened eyes [6]. However, thermally dimorphic fungi, although rarely, have also been reported as causative agents of FK [8]. Yeast ocular infections occur more frequently in temperate climates whereas filamentous fungal etiology is majorly documented in regions with tropical weather [20].

In this context, there are several epidemiological studies regarding the frequency of fungal keratitis, their related risk factors and the spectrum of etiological agents. In the era of the increasing number of immunocompromised populations and CLU, the prevalence of FK may be annually differing from country to country. Nevertheless, there is a lack of a comprehensive study comparing the prevalence rates of FK in different population-based studies, different countries from different continents, and the most frequent causative agents. Therefore, we aimed to systematically review data pertaining to studies concerning FK in the English language between 1990 and 2020 to provide contemporary insights into the epidemiology and causative agents of FK.

## 2. Methods

### 2.1 Database searching

The protocol of this study was registered in the International Prospective Register of Systematic Reviews (PROSPERO) with the ID number CRD42020188770 and the study was done according to the preferred reporting items for systematic reviews and meta-analyses (**supplementary file 1**) [21]. Relevant literatures were searched in Web of Science (ISI), PubMed, Scopus, and Google scholar using the main keywords “fungal keratitis”, “keratomycosis”, and “mycotic keratitis” and a set of other keywords, solely, and in combination (**supplementary file 2**). To ensure that the search captures all the relevant articles and because of usage of general phrases such as “infectious keratitis” or “microbial keratitis” that include a set of microorganisms i.e. bacteria, viruses, fungi, and amoebae; a wide search strategy was used. The search was limited to “article” as document type (whenever available), “English” as language, and “January 1, 1990 to the date of search (May 27, 2020)” as the publication date.

### 2.2 Study selection and quality assessment

The resulting articles in database searching were imported into an EndNote X9 software library for de-duplicating and title and abstract screening. After excluding irrelevant citations, full texts of citations were downloaded and checked for eligibility. All studies reporting data of prevalence and causative agents of fungal keratitis were eligible. Studies reporting animal or *ex vivo* models of keratitis, case reports and case series (without a denominator of the population that the cases had been diagnosed from), case-controls, cohorts, clinical trials, *in vitro* studies on virulence factors, antifungal susceptibility pattern of keratitis-isolated fungi but without data of prevalence, review articles, letters, and studies on therapy or keratitis caused by non-fungal microorganisms were excluded. Studies on specific populations e.g. specific age groups, those with specific surgical interventions, etc. were also excluded except for situations that several number of articles on the same patient group were available. In this case, the articles were included in the study but were analyzed separately. The quality of the relevant full texts was assessed using a modified version of the Newcastle-Ottawa Scale. All steps of screening and quality assessment were done by two independent researchers and in the case of inconsistency, a third researcher made the final decision.

### 2.3 Data extraction

Data of interest were the first author, year of publication, country, continent, the total number of studied patients, the number of fungal keratitis cases, frequency of yeast and mold pathogens, frequency of various fungal genera and species (if they were identified), frequency of mixed fungal and bacterial infections, gender and underlying conditions of confirmed patients (if available). These data were extracted by two independent researchers into a Microsoft Office Excel 2019 file.

### 2.4 Statistical analysis

Data were analyzed using Stata software version 14. To determine the heterogeneity, I^2^and Cochran Q test were used. In accordance with the Higgins classification approach, I^2^values above 0.7 were considered as high heterogeneity. In the presence of heterogeneity, a random effect model was used in calculations. The pooled prevalence with a 95% confidence interval (CI) was calculated using the “metaprop” command, and to estimate the pooled prevalence, we used the random-effect model. The exact method was used for calculating pooled estimates, variances, and their confidence intervals. We used Freeman-Tukey double arcsine transformation for variance stabilization.

The pooled prevalence of fungal keratitis and the pooled prevalence of yeast and mold keratitis and mixed fungal-bacterial infections were estimated. To determine the pooled prevalence of fungal keratitis in different countries, subgroup analysis was performed. The “metabias” command was used to check the publication bias, and if there was any publication bias, the prevalence rate was adjusted with the “metatrim” command using trim-and-fill method. The meta-regression analysis was used to examine the effect of the year of publication and sample size as factors affecting heterogeneity among studies. In all analyses, a significance level of 0.05 was considered.

## 3. Results

As presented in **Figure 1**, from 11235 articles retrieved in the primary search step, 169 met the inclusion criteria (**Supplementary file 3**). Results of their quality assessment are presented in **supplementary file 4**. These articles were divided into six groups, i.e. studies reporting data of fungal keratitis among (I) patients suspected of microbial keratitis (n=109), (II) suspected of fungal keratitis (n=13), (III) those with culture-confirmed microbial keratitis (n=10), (IV) contact lens wearers (n=6), (V) pediatric patients (n=8), and (VI) those who underwent keratoplasty (n=23), and analyzed separately. This practice was used for minimizing bias because the denominator was not identical in these groups. For instance, in group III, the denominator was very much smaller than other groups because it did not include patients who were clinically suspected and their etiology was rather non-infectious.

**Figure 1.**
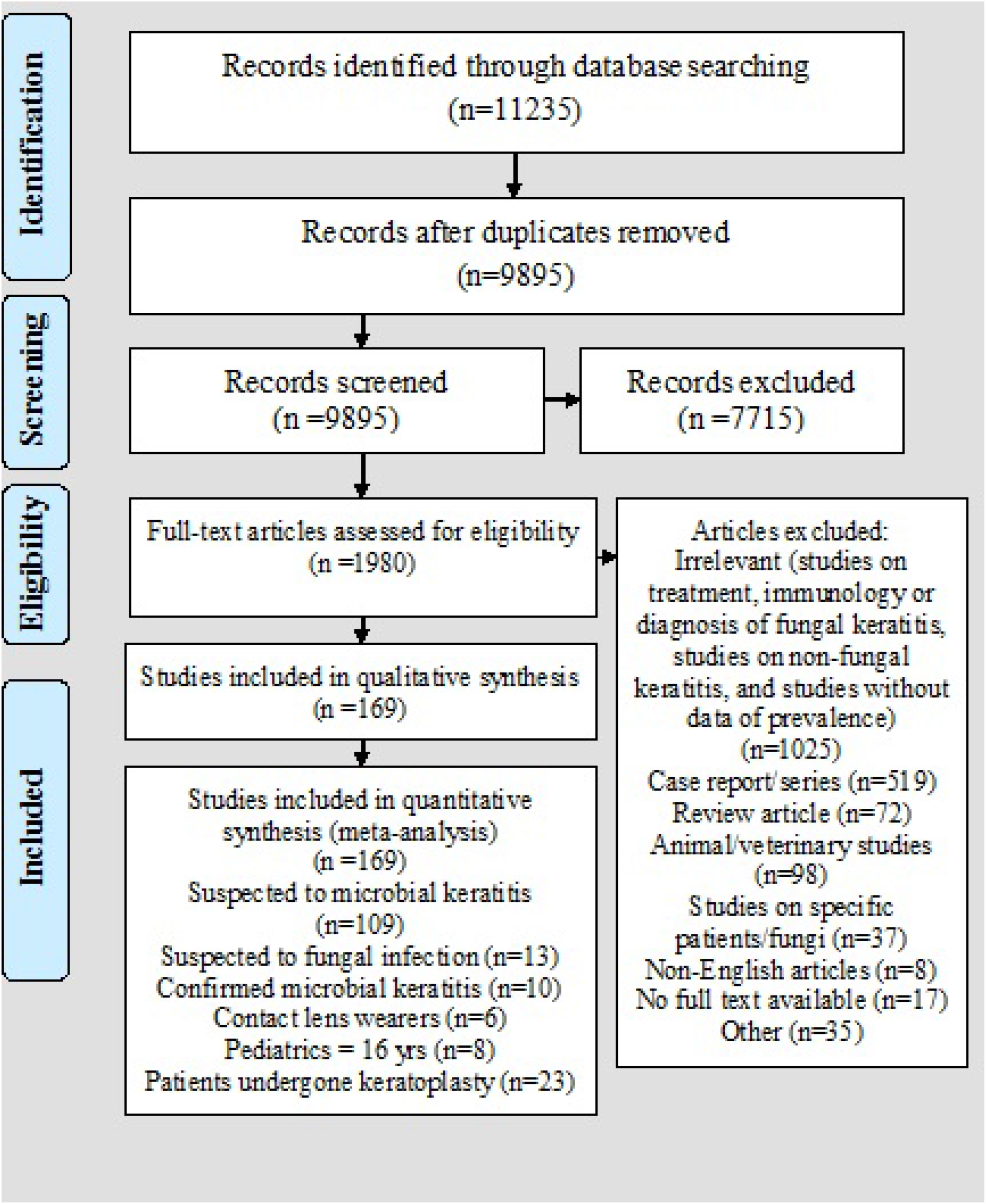
The PRISMA flow diagram of selecting studies reporting data on the prevalence of fungal keratitis between January 1, 1990 and May 27, 2020.

Regarding the origin of studies, from the 169 articles, 124 (72.94%) have been reported from Asia, 24 (14.12%) from America, 9 (5.29%) from Africa, 6 (5.29%) from Europe, and 4 (2.35%) from Oceania. In one article, two sets of data, one from India and one from Ghana have been reported, accordingly, they were treated as different studies in the calculation of origin of studies. Regarding the country of studies, the majority of studies have been reported from India (n=56, 32.94%), followed by China (n=16, 9.41%), USA (n=15, 8.82%), and Nepal (n=13, 7.65%). The distribution and frequency of studies from various countries are shown in **Table S1**.

### 3.1 Fungal keratitis among patients clinically suspected of microbial keratitis

In total, 109 articles were included in this group. Based on the analysis, the pooled prevalence of fungal keratitis among these patients was 23.64% (95% CI 20.39, 27.05) (**Figure 2**), and the prevalence of mold infections was found to be 87.01% (95% CI 83.31, 90.36) (**Figure S1**). There was no evidence of publication bias among these studies (**Figure S2**). Data of prevalence were available for 31 countries. According to the results of subgroup analysis which are shown in **Table 1**, the highest and the lowest prevalence has been reported from Paraguay (50.06%, 95%CI 35.11, 65.00) and Ireland (1.11, 95%CI 0.03, 6.04), respectively. Based on the results of meta-regression analysis, no significant change was noted in prevalence over the 30 years of study (*p*-value= 0.081) (**Figure S3**). There was also no association between the prevalence and the sample size studied in each report (*p*-value: 0.658) (**Figure S4**).

**Figure 2.**
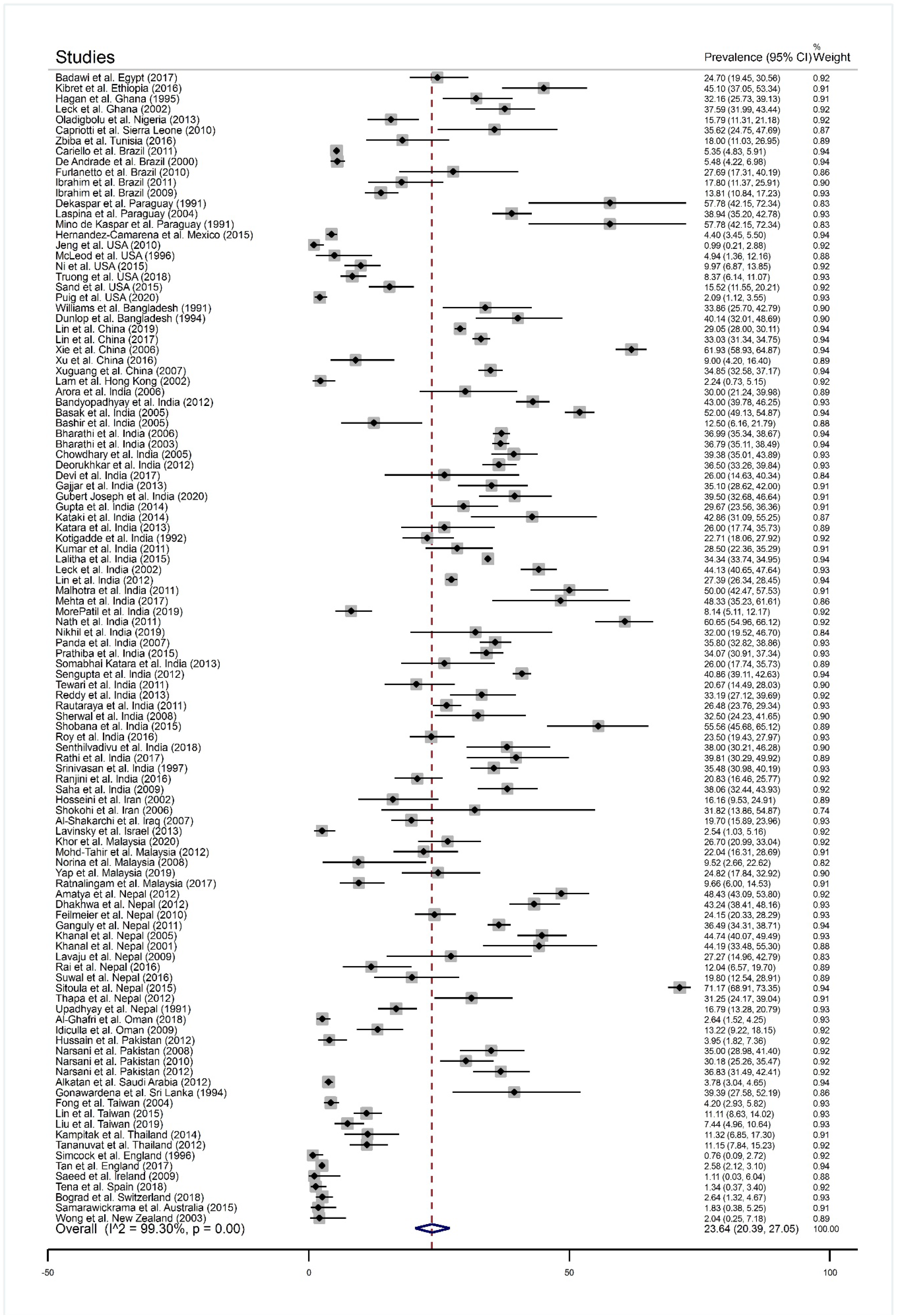
The forest plot of the prevalence of fungal keratitis among patients with a clinical suspicion of microbial keratitis based on the reported articles between January 1, 1990 and May 27, 2020.

**Table 1.** The pooled prevalence of fungal keratitis among patients clinically suspected of microbial keratitis in various countries (1990 up to May 27 2020)

| Country | Pooled prevalence | 95% confidence interval |
| --- | --- | --- |
| Paraguay | 50.06 | 35.11, 65.00 |
| Ethiopia | 45.10 | 37.05, 53.34 |
| Sri Lanka | 39.39 | 27.58, 52.19 |
| Bangladesh | 37.15 | 31.35, 42.95 |
| Sierra Leone | 35.62 | 24.75, 47.69 |
| Ghana | 35.35 | 31.16, 39.66 |
| Nepal | 34.42 | 23.38, 46.37 |
| India | 34.18 | 31.82, 36.58 |
| China | 33.18 | 23.64, 43.46 |
| Egypt | 24.70 | 19.45, 30.56 |
| Pakistan | 24.49 | 10.06, 42.71 |
| Iraq | 19.70 | 15.89, 23.96 |
| Malaysia | 18.44 | 11.48, 26.57 |
| Iran | 18.33 | 11.70, 25.96 |
| Tunisia | 18.00 | 11.03, 26.95 |
| Nigeria | 15.79 | 11.31, 21.18 |
| Brazil | 11.60 | 7.10, 17.01 |
| Thailand | 11.19 | 8.45, 14.25 |
| Taiwan | 7.30 | 3.54, 12.24 |
| USA | 6.06 | 2.33, 11.31 |
| Oman | 4.86 | 3.49, 6.43 |
| Mexico | 4.40 | 3.45, 5.50 |
| Saudi Arabia | 3.78 | 3.04, 4.65 |
| Switzerland | 2.64 | 1.32, 4.67 |
| Israel | 2.54 | 1.03, 5.16 |
| England | 2.37 | 1.94, 2.85 |
| Hong Kong | 2.24 | 0.73, 5.15 |
| New Zealand | 2.04 | 0.25, 7.18 |
| Australia | 1.83 | 0.38, 5.25 |
| Spain | 1.34 | 0.37, 3.40 |
| Ireland | 1.11 | 0.03, 6.04 |

From a total of 15295 fungal isolates, 13048 were identified. These isolates belonged to 63 distinct genera, and *Fusarium* species (n=5294, 40.57%) followed by *Aspergillus* species (n=4047, 31.02%) and *Candida* species (n=582, 4.46%) and these were the most common causes of the disease. As shown in **Table 2**, 628 of 5294 *Fusarium* isolates were identified at the species level and *Fusarium solani* which was the most common species. Similarly, 1405 of 4666 *Aspergillus* have been identified at the species level and *Aspergillus flavus* was the most common species. Among *Candida* isolates, 479 have been identified at the species level revealing *Candida albicans* as the most common species.

**Table 2.** The spectrum and frequency of various fungi isolated from patients with suspected microbial keratitis and identified at the genus/species level during 1990 to 2020 (13048 out of 15295 isolates have been identified)

| Fungus | Frequency (N=13048) | Percentage | Fungus | Frequency | Percentage |
| --- | --- | --- | --- | --- | --- |
| *Fusarium* | 5294 | 40.57 | *Trichophyton* | 21 | 0.16 |
| *Fusarium solani* | 361 | 2.77 | *Lasiodiplodia* | 21 | 0.16 |
| *Fusarium oxysporum* | 136 | 1.04 | *Fonsecaea* | 12 | 0.09 |
| *Fusarium moniliforme* | 116 | 0.89 | *Scopulariopsis* | 12 | 0.09 |
| *Fusarium nivale (Microdochium nivale)* | 11 | 0.08 | *Rhodotorula* | 10 | 0.08 |
| *Fusarium subglutinans* | 3 | 0.02 | *Cephalosporium* | 10 | 0.08 |
| *Fusarium verticillioides* | 1 | 0.01 | *Phialophora* | 8 | 0.06 |
| Unidentified species | 4666 | 35.76 | *Hormodendrum* | 7 | 0.05 |
| *Aspergillus* | 4047 | 31.02 | *Colletotrichum* | 6 | 0.05 |
| *Aspergillus flavus* | 562 | 4.31 | *Trichosporon* | 6 | 0.05 |
| *Aspergillus fumigatus* | 487 | 3.73 | *Verticillium* | 5 | 0.04 |
| *Aspergillus niger* | 330 | 2.53 | *Cylindrocarpon* | 5 | 0.04 |
| *Aspergillus terreus* | 17 | 0.13 | *Drechslera* | 4 | 0.03 |
| *Aspergillus nidulans* | 4 | 0.03 | *Chaetomium* | 4 | 0.03 |
| *Aspergillus oryzae* | 2 | 0.02 | *Exophiala* | 4 | 0.03 |
| *Aspergillus versicolor* | 2 | 0.02 | *Chrysosporium* | 4 | 0.03 |
| *Aspergillus tamarii* | 1 | 0.01 | *Trichoderma* | 3 | 0.02 |
| Unidentified species | 2642 | 20.25 | *Sepedonium* | 3 | 0.02 |
| *Candida* | 582 | 4.46 | *Phoma* | 3 | 0.02 |
| *Candida albicans* | 395 | 3.03 | *Nigrospora* | 3 | 0.02 |
| *Candida parapsilosis* | 44 | 0.34 | *Syncephalastrum* | 3 | 0.02 |
| *Candida tropicalis* | 28 | 0.21 | *Geotrichum* | 2 | 0.02 |
| *Candida glabrata* | 4 | 0.03 | *Epicoccum* | 2 | 0.02 |
| *Candida dubliniensis* | 2 | 0.02 | *Dichotomophthoropsis* | 2 | 0.02 |
| *Candida rugosa* | 2 | 0.02 | *Curwlanum* | 2 | 0.02 |
| *Candida krusei* | 1 | 0.01 | *Dichotomophthoropsis* | 2 | 0.02 |
| *Candida pelliculosa (Pichia anomala)* | 1 | 0.01 | *Diplosporium* | 2 | 0.02 |
| *Candida utilis* | 1 | 0.01 | *Microsporium* | 2 | 0.02 |
| *Candida guilliermondii* | 1 | 0.01 | *Monilia* | 2 | 0.02 |
| Unidentified species | 103 | 0.79 | *Graphium* | 2 | 0.02 |
| *Curvularia* | 777 | 5.95 | *Monosporium* | 2 | 0.02 |
| *Penicillium* | 389 | 2.98 | *Ustilago* | 1 | 0.01 |
| *Helminthosporium* | 332 | 2.54 | *Absidia* | 1 | 0.01 |
| *Mucor* | 305 | 2.34 | *Madurella* | 1 | 0.01 |
| *Cladosporium* | 230 | 1.76 | *Phaeosiarra* | 1 | 0.01 |
| *Alternaria* | 168 | 1.29 | *Cephalophora* | 1 | 0.01 |
| *Bipolaris* | 152 | 1.16 | *Keratomyces ajilloi* | 1 | 0.01 |
| *Botryodiplodia* | 143 | 1.10 | *Diplodia* | 1 | 0.01 |
| *Acremonium* | 133 | 1.02 | *Gymnoascus* | 1 | 0.01 |
| *Exserohilum* | 101 | 0.77 | *Epidermophyton* | 1 | 0.01 |
| *Scedosporium* | 56 | 0.43 | *Sporothrix* | 1 | 0.01 |
| *Aureobasidium* | 43 | 0.33 | *Cladophialophora* | 1 | 0.01 |
| *Zymoid epiphyte* | 41 | 0.31 | *Scytalidium* | 1 | 0.01 |
| *Rhizopus* | 35 | 0.27 | *Arthrographis* | 1 | 0.01 |
| *Paecilomyces* | 32 | 0.25 | *Cryptococcus* | 1 | 0.01 |

### 3.2 Fungal keratitis among patients with culture-confirmed microbial keratitis

From 169 articles, 10 met the inclusion criteria to be categorized in this group. These articles studied patients with a positive microbial culture. Accordingly, there was no patient free of microbial keratitis in this group. The pooled prevalence of fungal keratitis in this group was 17.89% (95% CI 6.96, 32.42) (**Figure S5**), of them, 76.63% (95% CI 53.16, 94.16) were due to molds (**Figure S6**).

From a total of 2016 fungal isolates from these patients, 1636 isolates were identified, mainly at the genus level. As shown in **Table 3**, the identified isolates belonged to 27 distinct genera, and *Fusarium* species (n=771, 47.13%) were the most common causes of disease followed by members of *Aspergillus* (n=584, 35.70%), *Candida* (n=82, 5.01%), and *Curvularia* (n=52, 3.18%).

**Table 3.** The spectrum and frequency of various fungi isolated from patients with confirmed microbial keratitis and identified at the genus level during 1990 to 2020 (1636 out of 2016 isolates have been identified)

| Fungus | Frequency (N=1636) | Percentage | Fungus | Frequency | Percentage |
| --- | --- | --- | --- | --- | --- |
| *Fusarium* | 771 | 47.13 | *Chrysosporium* | 6 | 0.37 |
| *Aspergillus* | 584 | 35.70 | *Cryptococcus* | 3 | 0.18 |
| *Candida* | 82 | 5.01 | *Rhizopus* | 2 | 0.12 |
| *Curvularia* | 52 | 3.18 | *Scytalidium* | 2 | 0.12 |
| *Acremonium* | 21 | 1.28 | *Chaetomium* | 2 | 0.12 |
| *Trichophyton* | 20 | 1.22 | *Exophiala* | 2 | 0.12 |
| *Bipolaris* | 16 | 0.98 | *Rhodotorula* | 2 | 0.12 |
| *Penicillium* | 11 | 0.67 | *Scopulariopsis* | 1 | 0.06 |
| *Alternaria* | 11 | 0.67 | *Beauveria* | 1 | 0.06 |
| *Exserohilum* | 11 | 0.67 | *Colletotrichum* | 1 | 0.06 |
| *Scedosporium* | 10 | 0.61 | *Epicoccum* | 1 | 0.06 |
| *Cladosporium* | 8 | 0.49 | *Chrysonilia* | 1 | 0.06 |
| *Lasioidiplodia* | 7 | 0.43 | *Fonsecaea* | 1 | 0.06 |
| *Paecilomyces* | 7 | 0.43 |  |  |  |

### 3.3 Fungal keratitis among patients clinically suspected of fungal keratitis

In general, 13 articles were categorized in this group. These articles included studies conducted on patients with a clinical suspicion of fungal keratitis. Thus, those with a clinical suspension of other types of microbial keratitis have been excluded. Based on the meta-analysis, the pooled estimated prevalence of fungal keratitis among these patients was 43.01% (95% CI 30.88, 55.59) (**Figure S7**), and in total, 91.76% (95% CI 87.34, 95.39) were due to molds (**Figure S8**).

From a total of 5557 isolates from these patients, 5245 were identified. As shown in **Table 4**, these isolates belonged to 42 distinct genera. *Aspergillus* species (n=1712, 32.64%) were the most common causes of the disease followed by *Fusarium* species. (n=1543, 29.42%), *Curvularia species* (n=480, 9015), and *Alternaria* species (n=478, 9.11%). Two isolates of thermally dimorphic fungi, i.e. *Blastomyces* and *Sporothrix* have also been recovered from these patients.

**Table 4.** The spectrum and frequency of various fungi isolated from patients with clinically suspected fungal keratitis and identified at the genus level during 1990 to 2020 (5245 out of 5557 isolates have been identified)

| Fungus | Frequency (N=5245) | Percentage | Fungus | Frequency (N=5245) | Percentage |
| --- | --- | --- | --- | --- | --- |
| *Aspergillus* | 1712 | 32.64 | *Absidia* | 2 | 0.04 |
| *Fusarium* | 1543 | 29.42 | *Exophiala* | 2 | 0.04 |
| *Curvularia* | 480 | 9.15 | *Scytalidium* | 2 | 0.04 |
| *Alternaria* | 478 | 9.11 | *Epicoccum* | 2 | 0.04 |
| *Candida* | 283 | 5.40 | *Acrophialophora* | 2 | 0.04 |
| *Bipolaris* | 255 | 4.86 | *Drechslera* | 2 | 0.04 |
| *Penicillium* | 197 | 3.76 | *Trichosporon* | 2 | 0.04 |
| *Acremonium* | 89 | 1.70 | *Scopulariopsis* | 1 | 0.02 |
| *Rhizopus* | 43 | 0.82 | *Cylindrocarpon* | 1 | 0.02 |
| *Paecilomyces* | 41 | 0.78 | *Neurospora* | 1 | 0.02 |
| *Rhodotorula* | 22 | 0.42 | *Lasiodiplodia* | 1 | 0.02 |
| *Scedosporium* | 16 | 0.31 | *Beauveria* | 1 | 0.02 |
| *Exserohilum* | 15 | 0.29 | *Phialophora* | 1 | 0.02 |
| *Cladosporium* | 12 | 0.23 | *Rhinocladiella* | 1 | 0.02 |
| *Mucor* | 7 | 0.13 | *Nigrospora* | 1 | 0.02 |
| *Fonsecaea* | 6 | 0.11 | *Papulaspora* | 1 | 0.02 |
| *Aureobasidium* | 5 | 0.09 | *Sarcinomyces* | 1 | 0.02 |
| *Chaetomium* | 4 | 0.08 | *Trichoderma* | 1 | 0.02 |
| *Trichophyton* | 4 | 0.08 | *Microsporum* | 1 | 0.02 |
| *Colletotrichum* | 3 | 0.06 | *Blastomyces* | 1 | 0.02 |
| *Chrysosporium* | 2 | 0.04 | *Sporothrix* | 1 | 0.02 |

### 3.4 Fungal keratitis among pediatric patients

Studies categorized in this group have been done on patients aged ≤16 years, except for 1 study which was on patients ≤15 years. Using this criterion, 8 articles were identified. The pooled estimated prevalence of fungal keratitis among these patients was 14.88% (95% CI 6.87, 25.11) (**Figure S9**) and molds accounted for 95.30% (95% CI 84.10, 100.00) of cases (**Figure S10**). Totally, 163 isolates were recovered from these patients, of them, 141 were identified (**Table 5**). These isolates belonged to 7 genera and *Fusarium* species (n=88, 62.41%) followed by *Aspergillus* species (n=31, 21.99%) were the most common causes.

**Table 5.** The spectrum and frequency of various fungi isolated from pediatric patients with fungal keratitis and identified at the genus level during 1990 to 2020 (141 out of 163 isolates have been identified)

| Fungus | Frequency (N=141) | Percentage |
| --- | --- | --- |
| *Fusarium* | 88 | 62.41 |
| *Aspergillus* | 31 | 21.99 |
| *Candida* | 16 | 11.35 |
| *Curvularia* | 3 | 2.13 |
| *Acremonium* | 1 | 0.71 |
| *Alternaria* | 1 | 0.71 |
| *Bipolaris* | 1 | 0.71 |

### 3.5 Fungal keratitis among contact lens wearers

Studies categorized in this group were those that reported the prevalence of fungal keratitis among non-therapeutic contact lens wearers. Using the inclusion criteria, 6 articles were identified, and based on the meta-analysis, the pooled estimated prevalence of fungal keratitis among them was 18.05% (95% CI 1.04, 46.91) (**Figure S11**). Almost all the cases (37 out o 38) were due to molds and the pooled estimated prevalence of mold infections was 100% (95% CI 94.30, 100) (**Figure S12**).

From 38 fungal isolates, 35 were identified as *Fusarium* species (n=31, 88.57%), *Aspergillus* species (n=3, 7.89%), and *Acremonium* species (n=1, 2.63%).

### 3.6 Fungal keratitis among post- keratoplasty patients

Generally, 23 articles met the inclusion criteria that were focused on patients who had undergone keratoplasty. We divided these articles into 2 groups, 7 articles that provided data on patients who had undergone keratoplasty due to a variety of indications except infective keratitis, and 16 articles that provided data on patients who had undergone keratoplasty due to infective keratitis. Such was done because the denominator of the latter group was smaller than that of the former. The pooled prevalence of fungal keratitis (reinfection or recurrence) was 0.05% (95% CI 0.00, 0.14) and 8.57% (95% CI 3.89, 14.62) in these groups, respectively (**Figure S13**). In general, the prevalence of yeast was higher among these patients with a pooled value of 51.80% (95% CI 14.41, 88.30) (**Figure S14**).

Data of the causative fungi were available only in 13 articles. From the total of 379 isolates, 294 were identified. As shown in **Table 6**, these isolates belonged to 13 distinct genera, including one isolate of *Pythium*, a member of Oomycota. *Fusarium* species (n=124, 42.18%) were the dominant cause followed by *Aspergillus* and *Candida* species. Among 27 isolates of *Aspergillus* that were identified to the species level, 22, 3, and 2 isolates were found to be *A. flavus*, *A. niger*, and *A. fumigatus*, respectively.

**Table 6.** The spectrum and frequency of various fungi isolated from keratoplasty patients with fungal keratitis and identified at the genus level during 1990 to 2020 (294 out of 379 isolates have been identified)

| Fungus | Frequency (N=294) | Percentage |
| --- | --- | --- |
| *Fusarium* | 124 | 42.18 |
| *Aspergillus* | 81 | 27.55 |
| *Candida* | 60 | 20.41 |
| *Curvularia* | 6 | 2.04 |
| *Alternaria* | 5 | 1.70 |
| *Acremonium* | 4 | 1.36 |
| *Penicillium* | 3 | 1.02 |
| *Pythium* | 3 | 1.02 |
| *Scedosporium* | 2 | 0.68 |
| *Bipolaris* | 2 | 0.68 |
| *Paecilomyces* | 2 | 0.68 |
| *Trichosporon* | 1 | 0.34 |
| *Wangiella* | 1 | 0.34 |

### 3.7 Mixed fungal and bacterial keratitis

The prevalence of mixed fungal and bacterial keratitis was calculated regardless of the grouping schedule. In this calculation, articles reporting data on keratoplasty recipients and patients with suspected fungal keratitis were excluded because the data of mixed infections were not available in almost all of these articles. From the remaining four article groups, data of mixed infections were extractable in 110 articles. Based on these analyses, the pooled prevalence of mixed infections was 9.29% (95% CI 6.52, 12.38).

## 4. Discussion

A systematic assessment of the 169 published literature from 36 countries (Table S1) revealed the geographic variation in the prevalence rates and predominant etiological genera of FK. A list of factors influence clinical outcomes of microbial keratitis and the epidemiological patterns differ from one country to the other, as well as different geographical regions of the same country [20]. In the present study, the majority of large scale investigations were reported from India, China, the USA, Nepal, Taiwan, and Brazil in descending rank, whereas the least studies were reported from Europe and Oceania continents which are principally because of the climatic conditions, non-agricultural activities resulting in low frequency of cases in these countries. Given that the highest pooled prevalence of FK were recorded in countries such as Paraguay, Ethiopia, Sri Lanka, and Bangladesh (Table 1) which had the lowest eligible studies included (Table S1); these prevalence rates might be less reliable than those documented in countries with high eligible studies such as India, China, USA, Nepal, Taiwan, and Brazil (Table 1 and Table S1). The general consensus is that FK is more frequent in developing countries within tropical and subtropical climate in comparison with developed countries with cold or temperate climate [5, 20]. Evidence is strong that the prevalence of FK is highly favored in areas with warm, humid climate and agricultural economy and its frequency has been estimated to range from 20 to 60% of all culture positive corneal infections in these regions [22]. The comprehensive data particularly the prevalence rates and predominant causal agents of FK in different regions and patient populations are indispensable to develop appropriate diagnostic and therapeutic strategies [5, 20]. There is still no systematic survey that assesses the rate of fungal keratitis in different population-based studies. This systematic review provides a global frequency of FK in different patient groups. The prevalence ratios given for FK depend on the settings and the studied population and were carefully compared, because of the different inclusion criteria used to select the studied subjects and the variations in sensitivity of the modalities used for FK diagnosis [22]. Depending on the populations examined, the prevalence of FK reported was up to 43% (95% CI 0.05-43%). The highest and lowest prevalence of FK were documented in patients with a clinical suspicion of keratomycosis and those who underwent keratoplasty respectively (43 % *vs*. 0.05%). The high prevalence evidenced in keratomycosis suspicion group in comparison with suspected-microbial keratitis patients (43% *vs*. 23.6%) was anticipated and is attributable to higher proportion of cases of clinically suspected FK than in studies which ocular infections might be caused by any of the fungal, bacterial, viral, amoebic, oomycete, or parasitic agents [23]. Similarly, former large-scale retrospective analyses investigated in India, Turkey, Paraguay, and Brazil reported much lower rates of FK (23%, 22.3%, 20.6 % and 5.3% respectively) in corneal cultures obtained from all patients with a suspicion of microbial keratitis [23–26]. However, regardless of variable frequency of FK in different geographical locations, the insensitivity of traditional microbiological methods (culture, smear stains) routinely used to diagnose microbial keratitis is accounted by the fact that many of the FK-affected people were resident of rural areas and might never refer to medical centers because of low income, cost of treatment, and long distance. It may also be due to the fact that most studies conducted at tertiary healthcare facilities, accepting more severe cases, resulting in overestimation of FK; bring together some drawbacks and limitation in assessing the real prevalence and incidence rate of FK in different studies.

Prevalence surveys done on patients with culture-confirmed microbial keratitis determined that 17.9% of microbial keratitis cases were caused by FK that was lower than both above mentioned groups. Considering the limitation of culture based methods [20], the lower rate of FK frequency documented in surveys including only culture positive specimens is expected. Although, culture methods remained the cornerstone of FK diagnosis in most studies [5] the isolation rates differed among different studies and the sensitivity of culture was as low as 50% [27]. It is also known that culture negative cases of FK may show fungal filaments in microscopic examination of the corneal scrapings and be diagnosed as FK [28]. Empirical treatment with topical antibiotics, topical anesthesia as thoughts to have antimicrobial effects, different methods of sample collections (swabbing *vs*. corneal scraping), the low quantity of specimens available for culture, refractive nature of fungi, and debatable adverse effect of transport devices or media on viability of microorganisms, are the possible confounders which may influence the results of culture method [5, 20, 27]. In the contact lens wearer group, 18% of the populations were rated to be infected with FK. In 25% to 40% of keratomycosis cases particularly those living in developed countries, contact lens wear is evidenced as a risk factor [12, 26]. Corneal defects, gradually caused by contact lens wear, poor hygiene practices such as overnight wear and ineffective or contaminated cleaning solution, have been increasingly associated with fungal keratitis [20, 22]. Contact lens-associated FK have majorly occurred in individuals with low socioeconomic status in which poor education about hygienic eye care and inadequate cleaning solution were accounted as attributable culprits [20, 22]. In the pediatric setting, the prevalence of FK was low (14.9%). Generally, men with agricultural and outdoor occupation and aged 20–50 years, form a greater proportion of the FK-affected population and are more susceptible than women to develop mycotic keratitis [5, 12, 20, 22, 26]. Nonetheless, children constitute about 4% of keratomycosis cases [22]. Children have minimal encounter with traumatizing agents (plant and animal sources), which are frequently associated with FK [22]. One study from southern California reported that pediatric keratitis composed 11% of all cases with microbial keratitis [29]. Another retrospective study from Taiwan reported that pediatric keratitis constituted 13.1% of all cases of infectious keratitis. Meanwhile, the frequency of pediatric fungal keratitis was as low as 6.4% among all culture positive pediatric microbial keratitis in Taiwanese children [30].

FK was rated to be 0.05% and 8.5% post-keratoplasty when the relevant studies were divided into two groups indicating the higher rate of prevalence in the population who underwent keratoplasty after infectious keratitis. For cornea maintained for more than 4 days in preservation medium, the relative risk of fungal keratitis was 3 times higher in comparison to bacterial infection, therefore supplementation of donor preservation media with an antifungal agent may be necessary [31]. Post-keratoplasty fungal keratitis in transplant recipients is majorly associated with infection of donor corneal tissue [32]. However, the higher rate of FK in populations who underwent keratoplasty after infectious keratitis was postulated to be unlinked to the donor corneal tissue but probably linked to the untreated or partially treated FK before the surgery that re-emerge after keratoplasty following immunosuppressive interventions. Although the infection is commonly rated to be rare [32], many ophthalmic surgeons believe that the incidence of post-keratoplasty fungal infections is rising. Fungal infection post endothelial keratoplasty (EK) is thought to be more frequent than penetrating keratoplasty (PK) [32]. The Eye Bank Association of America reviewed the frequency of keratoplasty-associated fungal infections and found that the development of fungal infections occurs in 0.052% of anterior lamellar keratoplasty procedures, 0.022% of EK procedures, and 0.012% of PK procedures [32].

Given the variations in antifungal susceptibility patterns of different fungal genera and even different species belonging to the same genus, definitive identification of the etiology to the species level is recommended [5, 9, 12, 20]. Nevertheless, in some studies, the nature of agents causing fungal infection was determined only by pathological sections [5]. On the other hand, the majority of early studies reporting the epidemiology of fungal keratitis have resorted to the identification of causative agents through culture-based morphologies and limited to the genus level [5, 20]. In the current review, we found the highest (94%) and lowest (77.5%) percentages of fungal identification (principally to the genus level) in studies done on cases with FK suspicion and keratoplasty patients, respectively. As a result, the identification of the investigated isolates to the species level has been provided in only a few reports. Therefore, the situation is not satisfactory. More so, the species identification is majorly achieved through morphology-based methods which may lead to a delayed or erroneous diagnosis in a significant number of cases. The subjective morphology based speciation is likely to be affected by the expertise of the investigator. Application of molecular based methods has been less frequent for species identification, with many causative agents, particularly the uncommon or previously unreported agents, as well as the non-sporulating molds remain unidentified [5, 20]. Since, a wide variety of fungal agents are known to be implicated in keratitis [5, 20, 22, 26], considerable mycological facilities, skills, and expertise are required for reliable identification of culture-positive cases and to rule out contaminants. The prevailing causal pathogens may vary in different geographical locations highlighting the need to know the local epidemiology [5, 20, 22, 26]. Similar to most of the studies across the globe [26, 33, 34], in almost all of our population based studies (except those done on patients with suspected FK), *Fusarium* species were the most predominant etiology of the disease followed by species belonging to *Aspergillus*, *Candida*, and *Curvularia* genera, which stand as the other frequent causes of FK (with a slight difference) in some studies. However, in tropical countries, southern United States, Mexico, Central America, South America, Africa, Middle East, China, India, and Southeast Asia, FK occurs mostly from filamentous fungi (particularly fusaria and aspergilli) and are frequently associated with plant material related corneal trauma, outdoor occupations and CLU [5, 12, 20]. Conversely, yeast associated mycotic keratitis (primarily due to *Candida* or *Cryptococcus*) were observed majorly in temperate countries, Europe and northern United States [5, 12, 20], and their infection is mostly linked to factors compromising the immunity of the eye such as local or systemic immunosuppressive agents used for corneal grafts and keratoplasty [5, 12, 20, 32]. Interestingly, in the present study, the highest percentage of *Candida*-associated keratitis (20.41%) was found in the group of patients undergoing keratoplasty (Table 6). *Fusarium* species are a serious threat to vision, especially for those wearing contact-lens. Consistently, the highest rate of *Fusarium* (88.5 % and 62.4%) was revealed in studies done on contact lens wearers and pediatric patients. The popularity of contact-lens wear is the leading factor predisposing these groups of patients to develop FK particularly with *Fusarium* etiology [27, 30]. The increasing prevalence of *Fusarium* keratitis was concurrently associated with a rising incidence of contact-lens wear [30, 35]. Interestingly, filamentous fungi (primarily *Fusarium* species) were almost sole agents causing keratitis in the contact lens wearers group in the current study. A similar resurgence of exclusive CLU-associated *Fusarium* keratitis was noted in literature from Hong Kong, Singapore, and United States [35–37]. CLU has evolved as an important risk factor for *Fusarium* keratitis [36].

Other than filamentous and yeast fungi, the dimorphic fungi are the third group scarcely reported as causal agents of FK [12]. In our review, two cases of FK due to *Blastomyces* and *Sporothrix* were observed among a wide variety of fungal agents causing keratomycosis in a population with suspected FK. Identification of the rarely reported fungi in this group of studies may be due to increasing utilization of molecular-based methods for fungal identification and better mycological skill and expertise of the researchers conducting these studies. In absence of application of molecular techniques, in some of the groups included in this study, the positive fungal cultures may have remained unidentified.

## 5. Conclusion

This review has illustrated the pooled prevalence of FK in different patient groups. The highest prevalence was demonstrated in the group of studies done on patients with suspected FK. Negative results of either microscopy or culture or both, in some corneal specimens obtained from patients with high clinical suspicion of FK, may affect the true rate of FK. This study suggests that this can be minimized if a high quality specimen from the clinic would be obtained and a trained and dedicated mycological service could be employed for diagnosis. Epidemiological variations within different countries are seen partly because of climatic situation and more so due to occupation of the population. Filamentous fungi such as *Fusarium* and *Aspergillus* continue to be the most frequently encountered genera in mycotic keratitis, and tend to be more predominant in traumatized eyes, CLU and pediatric groups. The highest rate of *Candida* species was recorded in patients with keratoplasty. Our data showed that the majority of the studies have used a culture-based method for the identification of causal agents up to genus-level and PCR-based identification molecular methods have been infrequently employed. As a result, species-specific therapy is hampered, particularly in the cases of less susceptible or resistant species of fungi.

## 6. Acknowledgments

## Conflicts of Interest

Nothing to declare.

## Funding

The authors declare that they have not received any financial support for this work.

---

**Figure S1.** The forest plot of the prevalence of yeast and mould keratitis among patients with a clinical suspicion of microbial keratitis based on the reported articles between January 1, 1990 and May 27, 2020

**Figure S2.** The funnel plot of available studies reporting data on the prevalence of fungal keratitis among patients with a clinical suspicion of microbial keratitis between January 1, 1990 and May 27, 2020. (each circle is representative of one study).

**Figure S3.** Univariate meta-regression analysis of the association between the year of publication and the heterogeneity. Each circle is representative of one study and its size shows the weight of observed effect sizes. The 95% confidence intervals of the regression line are shown by Error bars.

**Figure S4.** Univariate meta-regression analysis of the association between the sample size and the heterogeneity. Each circle is representative of one study and its size shows the weight of observed effect sizes. The 95% confidence intervals of the regression line are shown by Error bars.

**Figure S5.** The forest plot of the prevalence of fungal keratitis among patients with culture-confirmed microbial keratitis based on the reported articles between January 1, 1990 and May 27, 2020.

**Figure S6.** The forest plot of the prevalence of yeast and mould keratitis among patients with culture-confirmed microbial keratitis based on the reported articles between January 1, 1990 and May 27, 2020

**Figure S7.** The forest plot of the prevalence of fungal keratitis among patients with clinical suspicion of fungal keratitis based on the reported articles between January 1, 1990 and May 27, 2020.

**Figure S8.** The forest plot of the prevalence of yeast and mould keratitis among patients with clinical suspicion of fungal keratitis based on the reported articles between January 1, 1990 and May 27, 2020

**Figure S9.** The forest plot of the prevalence of fungal keratitis among pediatric patients based on the reported articles between January 1, 1990 and May 27, 2020.

**Figure S10.** The forest plot of the prevalence of yeast and mould keratitis among pediatric patients based on the reported articles between January 1, 1990 and May 27, 2020

**Figure S11.** The forest plot of the prevalence of fungal keratitis among patients with contact lens wearers based on the reported articles between January 1, 1990 and May 27, 2020.

**Figure S12.** The forest plot of the prevalence of yeast and mould keratitis among contact lens wearers based on the reported articles between January 1, 1990 and May 27, 2020

**Figure S13.** The forest plot of the prevalence of fungal keratitis among patients undergone keratoplasty based on the reported articles between January 1, 1990 and May 27, 2020.

**Figure S14.** The forest plot of the prevalence of yeast and mould keratitis among patients undergone keratoplasty based on the reported articles between January 1, 1990 and May 27, 2020

